# Collective pooling of foraging information in animal fission-fusion dynamics

**DOI:** 10.1101/2023.06.16.545019

**Authors:** Gabriel Ramos-Fernandez, Sandra E. Smith Aguilar

## Abstract

In animal species with fission-fusion dynamics, individuals can split from or follow others during collective movements. In spider monkeys (*Ateles geoffroyi*) this decision depends in part on the information they have about the location of available feeding trees. Foraging widely and continuously splitting and joining from others, individuals could be pooling their partial information such that the group as a whole has a more complete picture of a heterogeneous foraging environment. Here we use individual utilization areas over a realistic foraging landscape to infer the sets of potentially known trees by each individual. Then we measure the spatial entropy of these areas, considering tree species diversity and spatial distribution. We measure how complementary pairs of areas are, by decomposing the spatial entropy into redundant and unique components. We find that the areas uniquely known by each pair member still contain considerable amounts of information, but there is also a high redundancy in the information that a pair has about the foraging landscape. The networks joining individuals based on the unique information components seem to be structured efficiently for information transmission. Distributed foraging in fission-fusion dynamics would be an example of adaptive pooling of information and thus, collective intelligence.

## 2 Introduction

Sharing information about resource availability among group members has long been considered as one of the benefits of living in groups. Whether this sharing occurs actively, through specific signals, or more indirectly, through following knowledgeable individuals, collective foraging seems to be based on information transfer between individuals [Kummer, 2006,Petit and Bon, 2010,King and Sueur, 2011].

In species with a high degree of fission-fusion dynamics [Aureli et al., 2008] collective foraging occurs in a particularly distributed fashion, both in space and time. In these species, individuals form temporary subgroups of varying sizes that spread over large areas, travelling independently of other subgroups, but also coming together often. While the group as a whole is seldom all together in the same location at the same time, by splitting and joining often, individuals coincide frequently in the same subgroup after foraging in different regions of their home range. A subgroup then typically contains individuals who have knowledge about different areas of their home range.

In previous work [Palacios-Romo et al., 2019], we have shown that information about newly discovered fruiting trees can spread in this way in groups of spider monkeys. Through focusing our observations on relevant trees during their whole fruiting period, we showed that individuals that were naïve about the presence of fruit in the tree tended to arrive with other group members that already knew about the fruiting tree. Using a random arrival null model, we also showed that the group as a whole finds out about a particular food resource in fewer visits than if each individual had to find it on its own [Palacios-Romo et al., 2019].

A potentially beneficial outcome of information sharing in fission-fusion dynamics is that sub-group size is locally adjusted to the abundance of food. This resource matching benefit of fission-fusion dynamics has been shown in various studies, through partial correlations between the size of fruiting trees and the number of individuals feeding there [Symington, 1988, Chapman et al., 1995] or through an analysis of the distance traveled after a fission [Asensio et al., 2009]. Also, agent-based models of collective foraging show that agents sharing information about the location of target sites can exploit a heterogeneous environment more efficiently than when they only use their individual knowledge and coincide with others in the same targets by chance [Falcón-Cortés et al., 2019]. In further simulation studies, we have used the networks of influence during fissions and fusions and shown that subgroup size, and the way it responds to the overall abundance of food in the environment, is indeed sensitive to the probability that individuals will follow others during fissions or avoid each other before a fusion [Ramos-Fernandez et al., 2020].

Here we propose that this distributed form of collective foraging based on information transfer is a form of collective cognition, *sensu* [Theiner et al., 2010]. We define cognition as solving a specific problem by processing information. In particular, the problem would be how to find resources in a patchy, dynamic environment, while the solution would be matching subgroup size or visiting rates to the current availability of food. We explore the processing of information underlying this solution, by drawing on the partial information decomposition framework laid out by [Williams and Beer, 2010] (see also [Lizier et al., 2018] and [Bettencourt, 2009]) to measure the extent to which individual estimations of a target distribution of resources are synergistic (*i*.*e*. minimally redundant, maximally complementary) and show that spider monkeys might indeed be adaptively processing information in a distributed fashion [Hutchins, 1995].

## 3 Methods

### 3.1 Data collection

Ranging and subgroup composition data was collected by experienced observers during 20 months between January 2013 and September 2014 in a habituated group of black handed black spider monkeys (*Ateles geoffroyi*) living around the Punta Laguna lake in the *Otoch Ma’ax yetel Kooh* protected area, in the Yucatan peninsula, Mexico. This group was the subject of a long-term, continuous study from 1997 to 2020 [Ramos-Fernández et al., 2018].

We analyzed data for 29 individuals, including 16 adults (> 9 years or once they had an offspring), 5 subadults (5-9 years) and 8 juveniles (> 3 years or once they had a younger sibling). Observations were conducted during 4-8 hour daily subgroup follows. Subgroup scans were performed every 20 minutes, during which the subgroup composition was noted, as well as the position using a hand-held global positioning system located roughly in the center of the subgroup.

Individual core areas corresponded to the 60% utilization distribution generated with the Local Convex Hull (a-LocoH) method developed by [Getz et al., 2007]. These areas were calculated with a minimum of 100 scan sample locations for each individual, for a given season (Figure 2 shows a map of individual core areas for the wet season of 2014). For more details on the calculation of individual core areas, see [Smith-Aguilar et al., 2016].

Fruit abundance and phenological data was collected in a 1-ha plot in the middle of the group’s core area, in which all trees (with a diameter at breast height larger than 10 cm) from the 15 most consumed species by the spider monkeys (for a total of 487 individual trees) were assessed for the presence of fruit every 2 weeks from September 2013 to September 2014, comprising 25 monitoring periods.

### 3.2 Data analysis

We adapt the partial information decomposition method developed by [Williams and Beer, 2010] (see also [Lizier et al., 2018] and [Bettencourt, 2009]) to the problem of spatial foraging as follows. We assume there is a two dimensional foraging landscape *X*, which would contain the set of available fruiting trees at any given season within the home range of a spider monkey group. Each individual in the group of size *N*, 1, 2, …*N*, would be knowledgeable about a subset of *X, Y*_1_, *Y*_2_, …, *Y*_*N*_ as a result of their foraging movements within a limited area within *X* during a certain time period. These individual areas are estimations of the group’s foraging landscape, which is covered to different extents by overlapping combinations of these areas (Figure 1).

**Figure 1.**
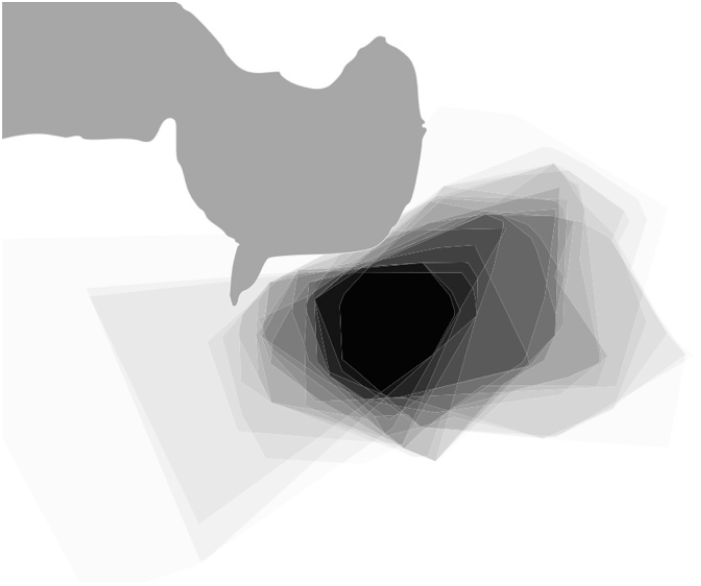
Example map of the 24 individual core areas for the wet season, 2014 overlaid on top of each other. The darker the shade of grey, the larger the number of individual core areas coinciding in a given point. The solid grey polygon on the top left represents the Punta Laguna lake as a reference. Although there is a high coincidence of core areas in the middle of the group’s range (solid black area), there is also a wide variation in the overall coverage of core areas, with large portions of the group’s range only covered by some individuals and not others.

The information decomposition method determines how synergistic (maximally complementary, minimally redundant) the estimations of the foraging landscape are. For illustration purposes, consider a group of size *N* =2 as in Figure 2. The information that *Y*_1_ and *Y*_2_ provide about *X* can be parsed into different components, as follows [Lizier et al., 2018]):

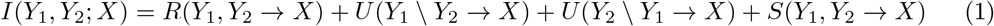

**Figure 2.**
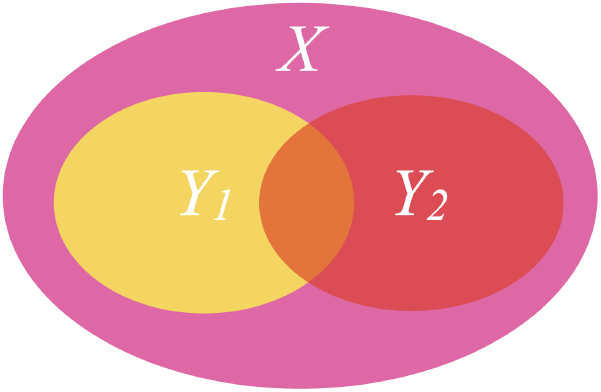
Diagram of the foraging landscape, *X* (pink area), and two individual estimations of *X, Y*_1_ (yellow) and *Y*_2_ (red). Information pooling will be synergistic to the extent that *Y*_1_ and *Y*_2_ are minimally redundant (orange area) and maximally complementary (high uniqueness, yellow and red areas; high synergy, sum of yellow and red areas) in their joint estimation of *X*.

Where *R*(*Y*_1_, *Y*_2_ →*X*) is the information about *X* provided *redundantly* by *Y*_1_ and *Y*_2_, *U* (*Y*_1_\*Y*_2_ →*X*) and *U* (*Y*_2_\*Y*_1_→*X*) are the *unique* contributions of *Y*_1_ and *Y*_2_, respectively, to *X*, and *S*(*Y*_1_, *Y*_2_ →*X*) is that component of *X* that can only be known by considering *Y*_1_ and *Y*_2_ together, or the *synergistic* information that they hold about *X*. Accordingly, each of the two individual estimations of *X* could be parsed into unique and redundant components, as follows:

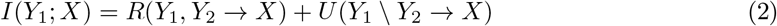

and

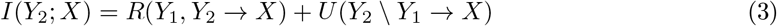

Where *R*(*Y*_1_, *Y*_2_ →*X*) corresponds to the redundant information that *Y*_1_ and *Y*_2_ contain about *X* (and would correspond to the orange area in Figure 2), both *U* terms refer to the information about *X* contained uniquely in either *Y*_1_ (yellow area in Figure 2) or *Y*_2_ (red area in Figure 2). *S*(*Y*_1_, *Y*_2_ → *X*) in equation 1, the synergistic information, would correspond to the union of the yellow, orange and red areas in Figure 2).

Translating this into spatial terms, we can think of the same areas in Figure 2 as containing trees of different species, which will have fruit at different times. The resulting distribution of available fruiting trees will be heterogeneous, both in space and time, and the group will need to continuously update its information about the location of available food. To the extent that *Y*_1_ and *Y*_2_ are complementary in their estimation of *X*, they will benefit by having a more complete and updated picture of their foraging environment.

In order to create a realistic foraging environment, we extrapolated each season’s fruiting tree abundance and distribution patterns from the 1-ha plot (see above) into a larger area which corresponded to the overlap of all individual core areas, which we assume is the foraging landscape, *X*, for a given season. Using the package *spatstat* [Baddeley et al., 2015] we created four different environments, corresponding to the dry and wet seasons of two different years for which we had data on space use. Because we only had data from the 1-ha plot on one dry season (2014), the extrapolation onto the foraging environment for the dry season in 2013 was done with the 1-ha plot data from the following season (see Supplementary Material).

The extrapolation of trees was done using a Poisson point process model assuming spatial inhomogeneity but with different intensities for each species that had fruit in a particular season. The assumption of spatial inhomogeneity is justified on the basis of a Ripley’s *K* test comparing the observed clustering of trees in the plot to the extrapolated environments (see Supplementary Material). This test uses the cumulative average number of points lying within a distance *r* of a typical data point (corrected for edge effects and normalized by intensity). Only in the 2013 wet season the parcel showed some evidence of clustering, with a few trees closer between 10-50m, but it showed no evidence of repulsion. In both 2014 seasons this does not seem to be the case, with a Poisson process assuming spatial inhomogeneity fitting the data well. In all cases, a Poisson process with different intensities for each species fit the observed data better than an alternative model that assumed only spatial inhomogeneity but no differences between species (see Supplementary Material). Figure 3 shows a map of each of these foraging landscapes, highlighting the differences in area shape and size, as well as a qualitative picture of the abundance and distribution of trees of different species.

**Figure 3.**
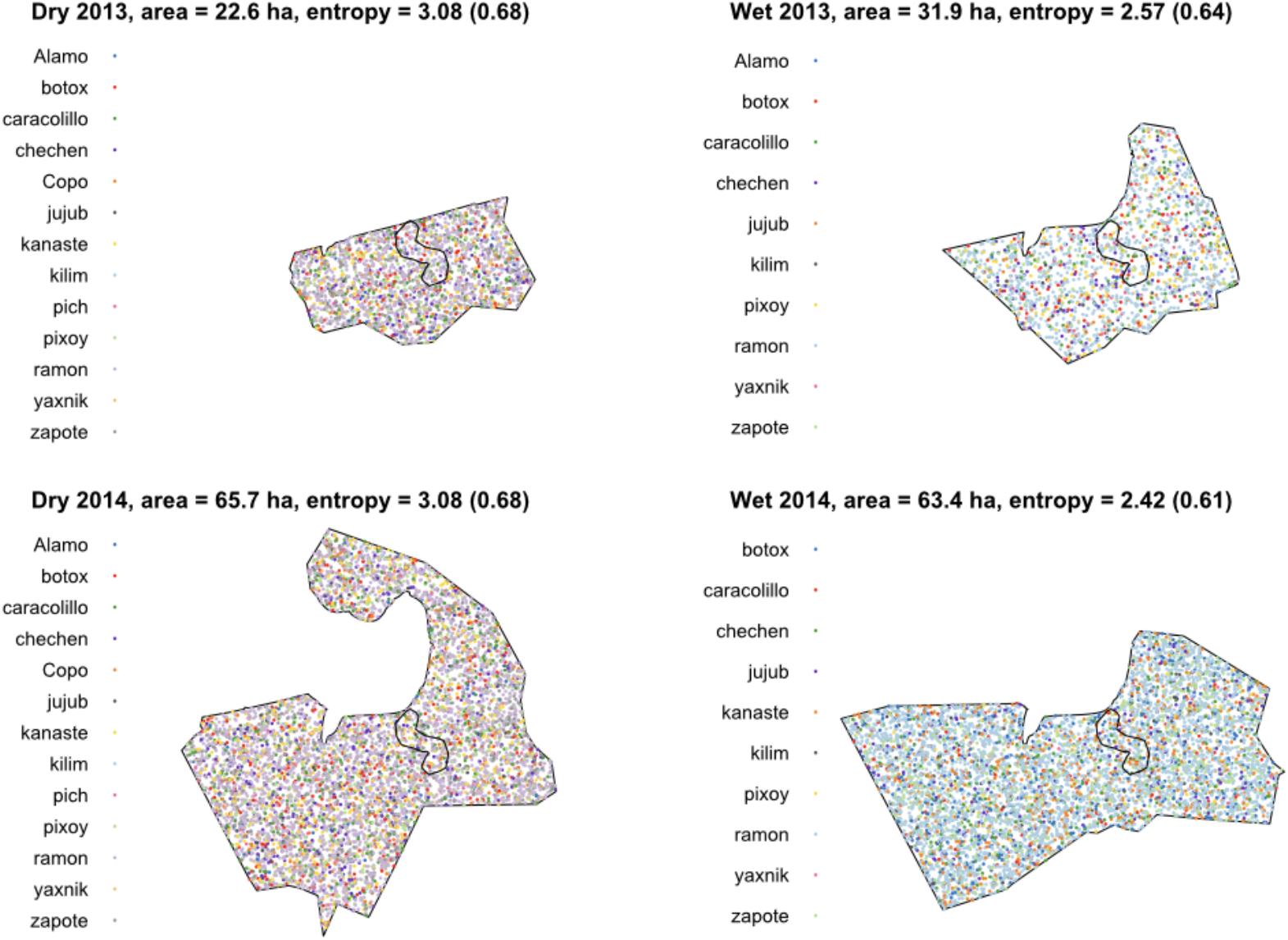
Maps of the extrapolated foraging landscapes for each of the four study seasons. Each point is a tree with available fruit colored according to species (common Maya names of the available species on the left key in each map), distributed according to a Poisson point process model based on the observed distribution in the 1-ha plot (wiggly polygon marked with a thick black line within each landscape). Also shown is the spatial entropy calculated for each foraging landscape, with the relative entropy, or the proportion of the maximum entropy possible, in parenthesis (see text for definitions). Total area for each landscape is given by the union of all individual corea areas (see Methods).

For measuring the information content of each foraging landscape *X* or core area *Y*_1_, *Y*_2_, …, *Y*_*N*_ we use the spatial entropy algorithm developed by [Altieri et al., 2021], available as a package for the R programming environment [R Core Team, 2021]. Briefly, this algorithm considers both the diversity of species within an area and the distances at which they can be found from one another. By evaluating a range of distances between pairs of trees, the heterogeneity of a foraging landscape (and thus its entropy) can be decomposed into a spatial component (i.e. at what distances are pairs of trees of similar species) and a species diversity component (i.e. which pairs of trees belong to the same species). A foraging landscape with little spatial autocorrelation and a high diversity of species would have a large entropy, while landscapes with less diversity and/or a high degree of clumpness or spatial autocorrelation by species would have a low entropy value. The Altieri spatial entropy (hereafter spatial entropy) can be expressed in bits or as the proportion of the maximum possible entropy an area could have, considering a maximum heterogeneity in the possible pairs of species found and their relative distances (hereafter relative spatial entropy). We assume that these information measures of a foraging landscape *X* in a given season are good estimations of the target knowledge that the spider monkey group as a whole should have, while for an individual core area they are good estimations of the knowledge that an individual has about the environment in a given season.

Given the above, if information pooling between the estimations *Y*_1_ and *Y*_2_ of a target distribution of resources *X* is synergistic, we predict the following:

1. Pairwise unions, *Y*_1_ ⋃*Y*_2_, should be large and have a similar entropy as the foraging landscape
2. Intersections, *Y*_1_ ⋂*Y*_2_, should be small and have a lower entropy than the foraging landscape
3. Set subtractions *Y*_1_\*Y*_2_, should be large and have a similar entropy as the foraging landscape
4. Because of the scarcity of resources compared to the wet season (see below), dry seasons should accentuate these effects

## 4 Results

The extrapolated foraging landscapes show a consistent seasonal variation in spatial entropy, which was higher in the dry seasons compared to their wet counterparts in both years (Figure 3). This consistent difference in entropy is in spite of the large difference in total area for both years (more than twice as large in 2014 than in 2013, for both seasons; Figure 3). The higher entropy in the dry seasons could be due, in part, to the higher number of species with fruit (13 vs. 10 in the wet seasons), but not exclusively, since the spatial entropy algorithm we have used takes into account not only the diversity but also the spatial distribution and abundance of the different species. It is important to note that a larger number of relevant species during the dry season does not necessarily imply a higher abundance of overall resources. The wet season contains higher overall abundances of fruit, but from fewer species [Smith-Aguilar et al., 2016].

The pairwise unions are much larger than the intersections and set subtractions and, as predicted, have a similar spatial entropy as the foraging landscape (Figure 4; area ANOVA test with surface type *F*_1,97_ = 559.85, *P <*0.001; year *F*_1,62_ = 387.44 *P <*0.001 and season *F*_1,62_ = 110.89, *P <*0.001 as factors; spatial entropy ANOVA test with surface type *F*_1,63_ = 35.13, *P <*0.001; year *F*_1,62_ = 28.4, *P <*0.001 and season *F*_1,62_ = 3198, *P <*0.001 as factors). This is important because the pairwise unions represent only a portion of the foraging landscape—few pairs of individuals have unions encompassing more than a half of the area of the foraging landscape. This implies that in terms of area, a complete picture of the foraging landscape is acquired by sets of individuals that are larger than a pair. However, the information content of these pairwise unions is comparable to that of the foraging landscape.

**Figure 4.**
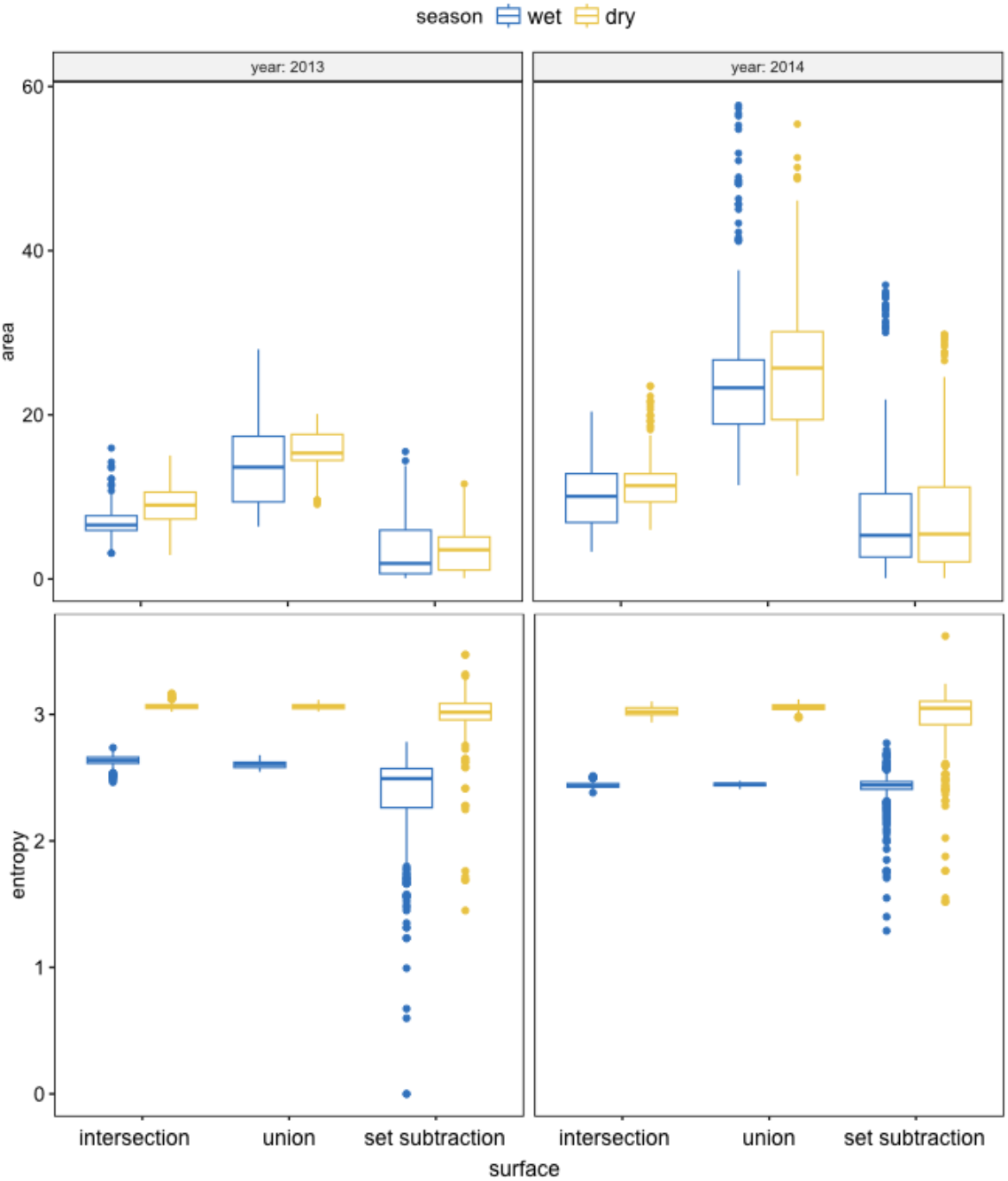
Area (upper panel) and spatial entropy (lower panel) for the intersections, unions and set subtractions of pairwise individual core areas. Each box represents the distribution of values of all pairwise combinations for each polygon type, during a given season and year.

The fact that intersections tend to be small implies low levels of redundancy in the information shared by pairs, although their spatial entropy is similar to the unions’ and also comparable to that of the foraging landscape as a whole, differing between years and seasons in the same way as unions did, being clearly higher during the dry seasons of both years. Finally, set substractions were smaller than intersections in both study years and across seasons. Set subtractions showed very variable levels of spatial entropy, which on average was lower than the unions’ and intersections’. This implies that, in general, the unique information is lower than the redundant information.

Even though they are smaller and have a lower average spatial entropy than the unions and intersections, set subtractions [*Y*_1_ *\ Y*_2_] clearly identify which areas are uniquely used by one member of the pair and not the other. This could reflect how information about resources is transferred between two individuals, as *Y*_1_ visits some areas that *Y*_2_ does not. At the same time, both individuals coincide in the intersection of their core areas, which could allow them the opportunity to share information about those uniquely known resources. Figure 5 shows an example of two partially overlapping core areas and how their set subtractions represent unique foraging information.

**Figure 5.**
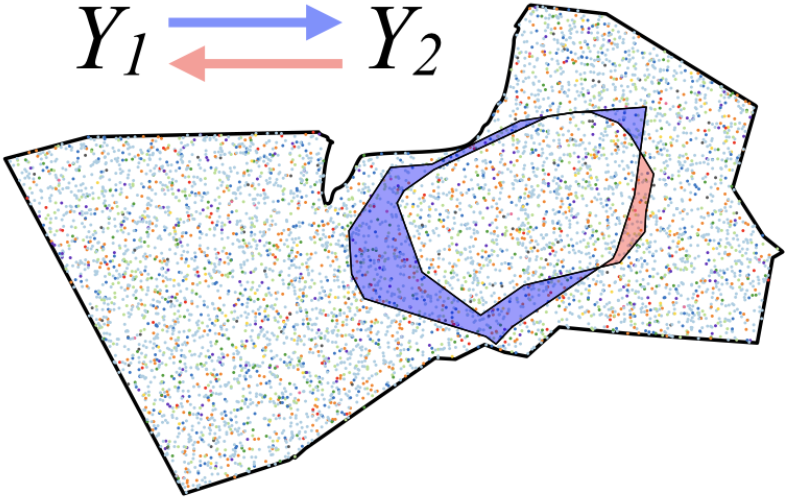
Example of two individual core areas overlapped inside the wet season 2014 foraging landscape. The blue area represents the set subtraction [*Y*_1_ *\ Y*_2_], or that area covered by *Y*_1_ but not by *Y*_2_, and the orange area the set subtraction [*Y*_2_ \ *Y*_1_], or the area covered by *Y*_2_ but not by *Y*_1_, during that season. This can be interpreted as the unique information that *Y*_1_ has that could be transmitted to *Y*_2_ (blue arrow), and the unique information that *Y*_2_ has that could be transmitted to *Y*_1_ (orange arrow).

As expected from the boxplots in Figure 4, the matrices of set subtractions show a wide range of variation in relative spatial entropy (Figure 6). These matrices are asymmetric, due to the fact that [*Y*_1_ \ *Y*_2_] ≠ [*Y*_2_ \ *Y*_1_]. In all seasons we find many absent values, due to those individual areas that lie completely within others and therefore have a null set subtraction. However, the fact that most values of relative spatial entropy lie above 0.6 implies that for most pairs, the unique information that *Y*_1_ has about *X* that *Y*_2_ does not have is still significant. There are some cases, especially in the dry seasons, where individuals may use unique areas not shared by another and that may contain 80-90% of the maximum possible information. We also find few balanced, bi-directional relationships where two individuals share a similar amount of information, suggesting that there is a directional flow of information from individuals that are more knowledgeable to those that are less.

**Figure 6.**
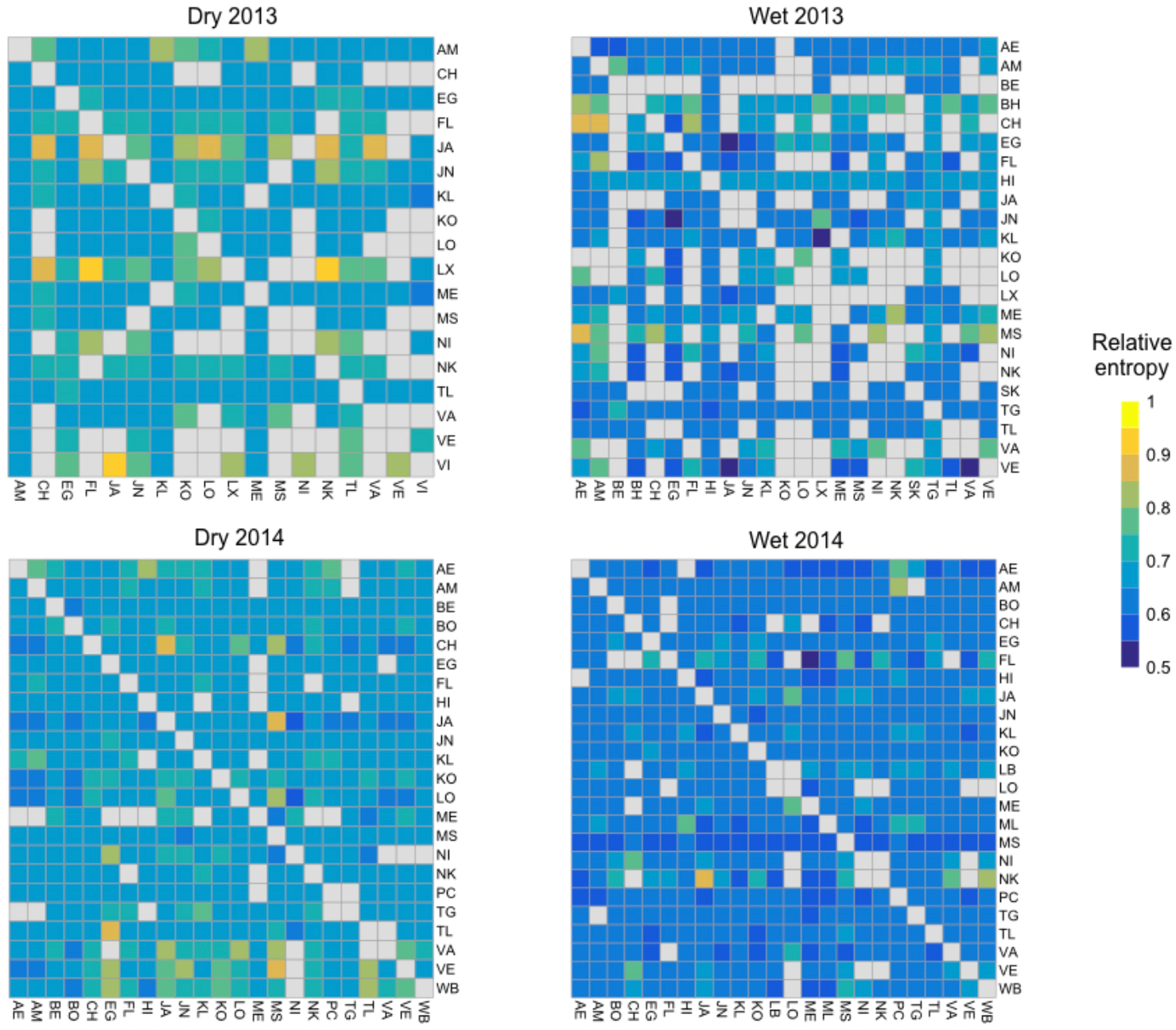
Heatmaps showing the spatial entropy of the set subtraction between pairs of individuals for the different seasons. In order to make them comparable between seasons, we show relative spatial entropy values, which reflect the proportion of the maximum possible entropy of a given area (see Methods). The entries in these asymmetric matrices can be read as the amount of information that an individual *Y*_1_, in rows, could be transferring to another individual *Y*_2_, in columns, by virtue of using an area that *Y*_2_ does not use, for a given season. Grey entries correspond to absent values of spatial entropy due to null set subtractions of individual areas that lie completely within another one. The code names of individuals label each row or column, with age and sex class as in Figure 7.

**Figure 7.**
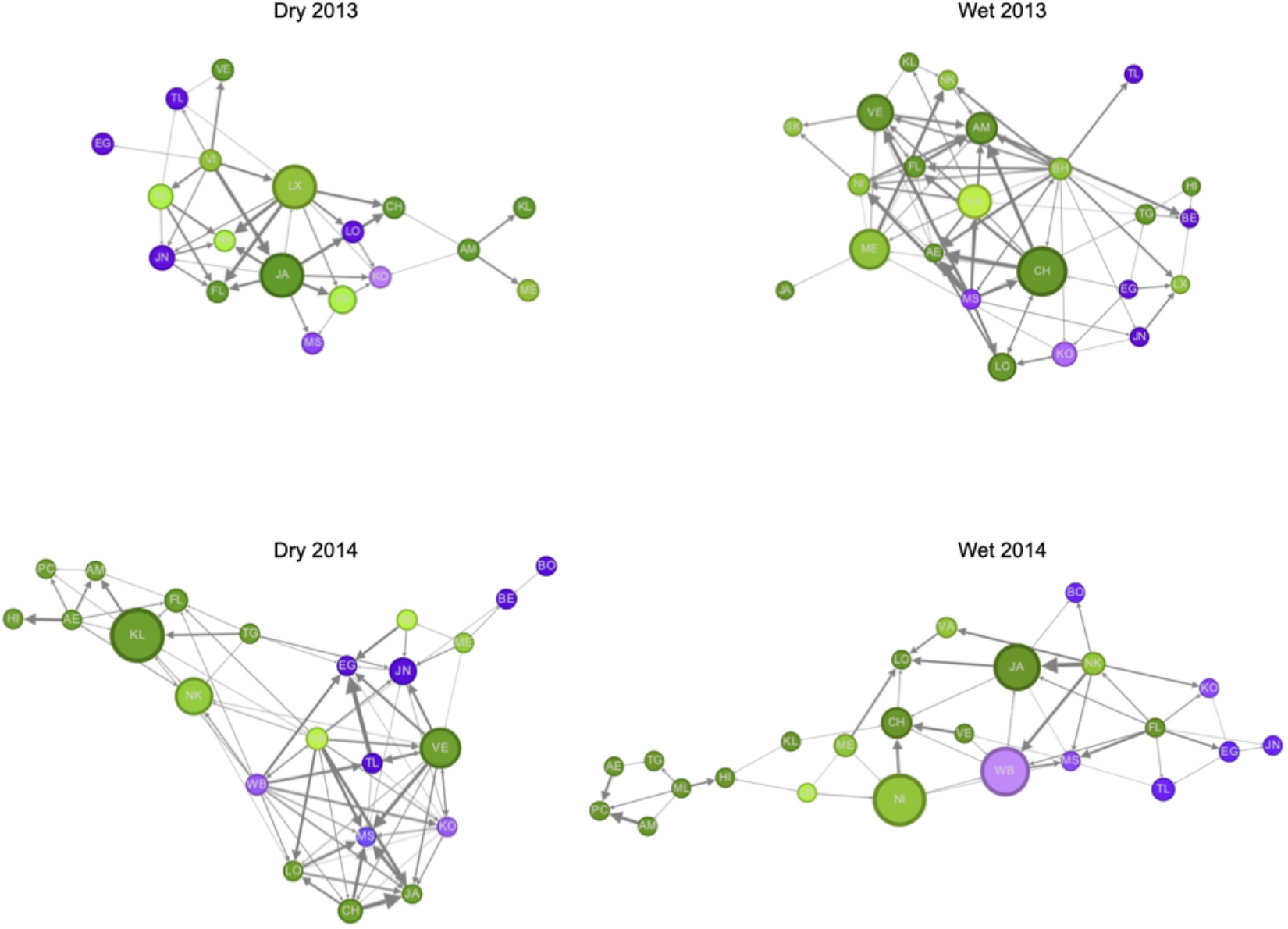
Networks of relative spatial entropy with the entries in the matrices in Figure 6 represented as directed links of proportional width. A link represents the relative spatial entropy in the set subtraction from one node *Y*_1_ to another *Y*_2_, or the amount of unique information that flows from *Y*_1_ to *Y*_2_. Nodes represent individuals of different age and sex classes (green for females, purple for males, tones decreasing in darkness from adults to subadults to juveniles). Link weights were filtered to show only those with relative entropies above 0.65, which was the value around which networks remained in a single connected component. Node size corresponds to the betweenness centrality within the filtered network.

A more detailed analysis of the information that *Y*_1_ has about *X* that *Y*_2_ does not possess is afforded by the directed, weighted network of set subtractions. In these networks, two individual nodes *Y*_1_ and *Y*_2_ are linked in proportion to the relative spatial entropy of the set subtraction *Y*_1_ \ *Y*_2_, or the information that *Y*_1_ has about *X* that *Y*_2_ does not have. These networks show the direction in which unique foraging information could be flowing between individuals in the group. They highlight the fact that, for the great majority of pairs of individuals, unique information seems to be flowing through other individuals. In other words, many individuals are distantly connected to others in the network and thus must obtain only partial foraging information from their immediate neighbors in the network. At the same time, some individuals seem to receive and give more information than others, and thus become more central in terms of their betweenness, or how often they lie in the shortest path between all other pairs of individuals. Clearly, adult females have a higher betweenness centrality than all other age and sex classes.

Finally, we consider the way in which overlaps of more than a pair of individual areas could lead to collective information pooling. For this, we identified the areas covered by 1-4 core areas and classified them as low overlap, then those covered by 5-15 core areas and classified as medium overlap areas, and finally those covered by 16 or more layers, which we considered high overlap areas. We measured the spatial entropy of these areas in each season and year combination (Figure 8). While showing the same pattern of dry seasons having a higher spatial entropy than wet seasons in both years, the low overlap areas, which presumably consist of those areas that are explored more occasionally when there is food scarcity, showed a higher spatial entropy in the dry season compared to the wet seasons, when they were consistently the areas with the lowest spatial entropy.

**Figure 8.**
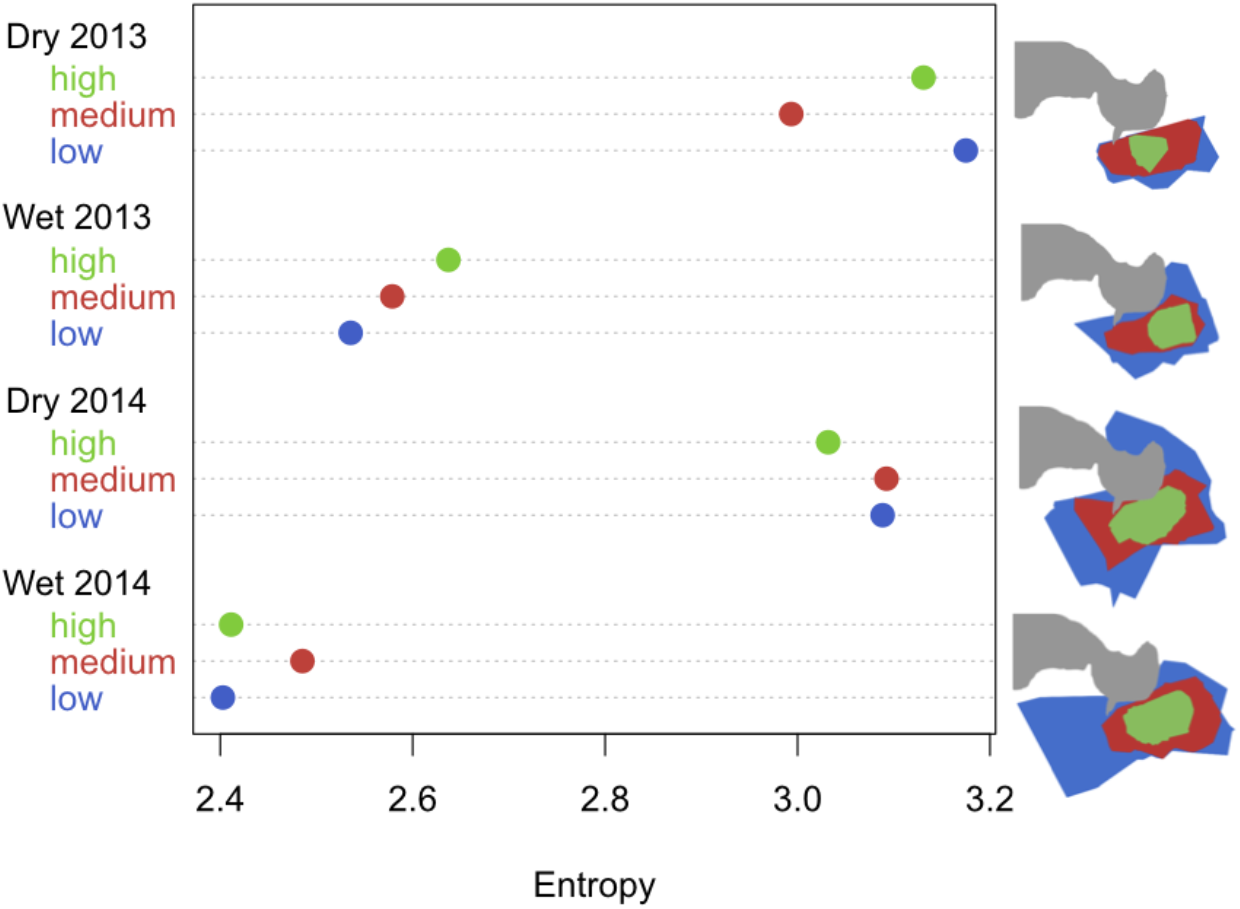
Values of spatial entropy for the areas with low overlap (1-4 individual core areas), medium overlap (5-15 individual core areas) or high overlap (16 and over individual core areas). Shown to the right are schematic maps of these areas for the different season and year combinations, with the lake area shown in grey. As also evident from Figure 1, regions of high overlap lie in the middle of the foraging landscape and are surrounded by medium and low overlap areas.

## 5 Discussion

We have used a spatial entropy analysis of individual space use areas to evaluate the way in which different individuals in a fluid society could be sharing information about a foraging landscape that is continuously changing. While we find a high degree of overlap between individual core areas and therefore a high redundancy in the information that each individual has about the foraging landscape, we also find areas that are uniquely known by individuals, with varying levels of spatial entropy. By constructing networks with links proportional to the information that is known by an individual but unknown by the other, we can obtain a picture of how foraging information could be flowing amongst group members. Going beyond the dyadic level, we also find that areas with less individual layers overlapping have higher spatial entropy during the dry season, when resources are scarcest, which suggests a more distributed exploration of a less abundant but more diverse foraging landscape. By sharing information that individuals know uniquely, spider monkey groups could be thought of as collectively intelligent [Hutchins, 1995, Theiner et al., 2010].

Early applications of information theory to collective animal foraging date from the work of [Haldane and Spurway, 1954] and [Wilson, 1962]. These works measured, for bees and ants, respectively, the information content of the message transmitted between individuals, based both on the variability of the message as well as on the behavior of its recipients. Here, we do not study a particular communicative signal which is supposed to be informing other group members about sources of food, even though some of spider monkeys’ vocalizations have been proposed to have that function [Ramos-Fernandez, 2008]. Rather, we assume that by following knowledgeable individuals, others can find food sources unknown to them, a process we showed could be at work in a previous study which used detailed observations of who visited newly found fruiting trees and with whom [Palacios-Romo et al., 2019]. Based on this finding, and noting the high degree of fission-fusion dynamics of the species, we assume that individuals could be sharing information about those fruiting trees they know uniquely during the time they spend in the same subgroup with other individuals that have been ranging elsewhere. The collective effect of this information sharing would be a more accurate picture of the foraging landscape than any individual could gain on its own. We use information theory to measure this knowledge sharing, particularly its unique and redundant components.

We find that our results on area and spatial entropy are related, with larger areas having higher spatial entropy. This is to be expected from the definition of spatial entropy we employ here, which considers the diversity of species as well as the distribution of distances at which pairs of different species can be found. The larger the area, more pairs from more different species can be found and thus the spatial entropy will increase. However, entropy and area do not necessarily speak of the same thing. Two areas of similar size can contain widely different sets of trees, of species that have very different abundance and spatial distribution patterns. The differences we find between the dry and wet seasons, for areas of similar size in each study year, are an example of this. But two individual core areas of similar size could also have different spatial entropy based on factors that are actually in the data like the areas’ shapes, for example.

We found that the clearest effect on spatial entropy is the season. Dry seasons present a more diverse, yet less abundant foraging landscape, which increases its spatial entropy and that of all the individual areas and their intersections and set subtractions. During these seasons, spider monkeys rely on more, less abundant species and thus exploit a more heterogenous environment. The study site has a hyperabundance of ramon (*Brosimum alicastrum* [Ramos-Fernández et al., 2018]), which when in fruit during the wet season would make it less valuable to share information about scarce sources of food. This would explain why in our analysis, areas with lower overlap, which presumably result from infrequent explorations of marginal areas of the core areas by a few individuals, have a higher spatial entropy during the dry season compared to areas with higher levels of overlap. Thus, during the dry seasons, when food is more scarce and dispersed and subgroups tend to be smaller and home ranges larger, those parts of the core areas of few individuals represent more information for the group. This would be in line with the idea of information pooling because in the face of scarcity, information on the few food resources available becomes more relevant than, for instance, in the wet season when resources are abundant.

We predicted that pairwise unions of individual areas would be larger than intersections and set subtractions, and that they would have a comparable information content as the foraging landscape. This was indeed the case, even when individual areas are considerably smaller than the foraging landscape. It is yet more interesting to compare the redundant unique components of unions, since their balance represents how complementary is the pooling of foraging information between pairs of individuals [Lizier et al., 2018]. We predicted that set subtractions would be large and have a similar spatial entropy as the foraging landscape, while intersections would be small and have a lower spatial entropy than the foraging landscape. We did not find this, as set subtractions had the lowest spatial entropy levels overall, while intersections were comparable to the unions. This would imply that pooling of information is not so complementary under these definitions. However, it should be noted that both unique and redundant areas both direct consequences of space use in a species with a high degree of fission-fusion dynamics: two individuals may be found in the same subgroup during part of the time, thus occupying the same areas, while they may spend time in different subgroups and potentially explore different parts of the foraging landscape [Smith-Aguilar et al., 2016]. Thus, even when there is a necessary degree of redundancy in how much information each individual contributes with to the common pool, the fact that there is unique information between all pairs, even when a complete picture of the foraging landscape can only be observed by looking at sets of individuals that are larger than a pair (see Figure 1, for example) implies some degree of complementarity in the information pooling by the group as a whole.

It is important to note that by analyzing core areas defined as 60% utilization distributions, it is possible that the regions of the set substraction *Y*_1_ \ *Y*_2_ areas we have assumed to be unknown by each member of the dyad could in fact be known if we looked at a wider utilization distribution such as 95% (frequently used to define home ranges [Laver and Kelly, 2008, Asensio et al., 2012]). We chose core areas because they tend to capture heavily used parts of the home range which concentrate high quality resources such as food sources and sleeping sites [Powell, 2000, Asensio et al., 2012]. Therefore, we assume that in core areas, individuals have a higher chance of knowing the available food sources than in less intensely used areas of their home range, where individuals would be more commonly naïve about trees with fruit. Therefore, when stating that the part of set substraction area *Y*_1_ \ *Y*_2_ corresponding to *Y*_1_ is unknown to *Y*_2_, it could be more accurately said that it will be more likely to have foraging information unknown by *Y*_2_ compared to *Y*_1_.

The directional, weighted networks we construct with the set subtractions, which we assume to represent the flow of information about uniquely known information between pairs, show that individuals can vary widely in their betweenness centrality, a measure of how relevant they are in the shortest paths between pairs of individuals. If an individual that receives information from another can in turn share it with a third, then these individuals with high betweenness centrality could be considered to be the most relevant for maintaining the flow of information about newly found sources of food in the group. At the same time, the fact that the networks are cohesive, with a single connected component, also speaks to the fact that even when not coinciding in the same social unit at the same time, spider monkeys do function as a group, even though sharing information about food sources in a manner that is displaced in time. The existence of a single connected component and some individuals with high centrality functioning as intermediaries between different regions of the network, could constitute an efficient network structure for information diffusion [Newman, 2003, Voelkl and Noë, 2010, Centola, 2022].

In addition, we can compare information-sharing networks in [Palacios-Romo et al., 2019] to those found here. In that study, we defined a link based on the frequency with which an individual that was naïve about a fruiting tree had arrived to it with another individual that in that moment was already knowledgeable about the same tree. In those networks, adult females seem to be more central and there seems to be more sex segregation, reflecting the more typical social behavior of the species [Fedigan and Baxter, 1984]. Here, the networks we have uncovered reflect who could potentially transfer more information to another because of their set subtractions. We find less sex segregation in these networks, with some adult females occupying a highly central position in some, but not all, the study seasons. Both networks could be reflecting the information sharing process, one at a local, patch level and the other at a foraging landscape level, and both could be complementary in providing the spider monkeys with a way to forage collectively in a closer to optimal way.

In conclusion, we have shown how a more accurate picture of a complex, dynamic foraging landscape could be obtained by the group when individuals pool unique information about available fruiting trees. We have done this through the application of an information-theoretic framework to quantify collective intelligence in a species with highly fluid grouping patterns. This would be an example of collective intelligence in natural conditions [Hutchins, 1995].

## 6 Acknowledgements

We thank Linda Altieri for her advise with the spatial entropy algorithm. Fieldwork was carried out under research permits DGVS00910/13 and DGVS02716/14 from the Ministry of Environment and Natural Resources of Mexico (SEMARNAT). Funding was provided by grants 157656, 207883 and CF-2019-263958 by the National Council for Science and Technology. We thank the Diverse Intelligences Summer Institute (DISI 2023) for the opportunity to discuss this work.

## Supplementary material for

Here we provide more details on the way we created a realistic foraging environment by extrapolating each season’s fruiting tree abundance and distribution patterns as measured in a 1-ha plot (see Methods) into the larger foraging landscape for each season (which corresponded to the overlap of all individual core areas). All analyses were performed using the *spatstat* package for *R* [R Core Team, 2021,Baddeley et al., 2015]. For two wet seasons and one dry one we had data on all trees with fruit and their location within the 1-ha plot. For the dry season in 2013 we only had observations of monkeys but no data on fruiting patterns. Therefore, we used the same model for the dry season in 2014 to extrapolate into the overlap of individual core areas in the dry season, 2013.

We assumed that the distribution and abundance of fruiting trees in the plot could be reasonably well described by Poisson point process model which assumed spatial inhomogeneity and a different intensity for each species. Spatial inhomogeneity implies that there is neither significant clustering nor repulsion between trees. This assumption was justified by means of a Ripley’s *K* test and a comparison of the Poisson model with and without different intensities for each species.

The Ripley’s *K* test compares the cumulative average number of trees lying at different distances from a typical data point with the theoretical expectation assuming a Poisson process with spatial inhomogeneity. Figure 1 shows the fit between the theoretical and observed curves, which is better for both 2014 seasons than for the wet season of 2013, when there is some evidence of a higher intensity of clustering.

**Figure 1.**
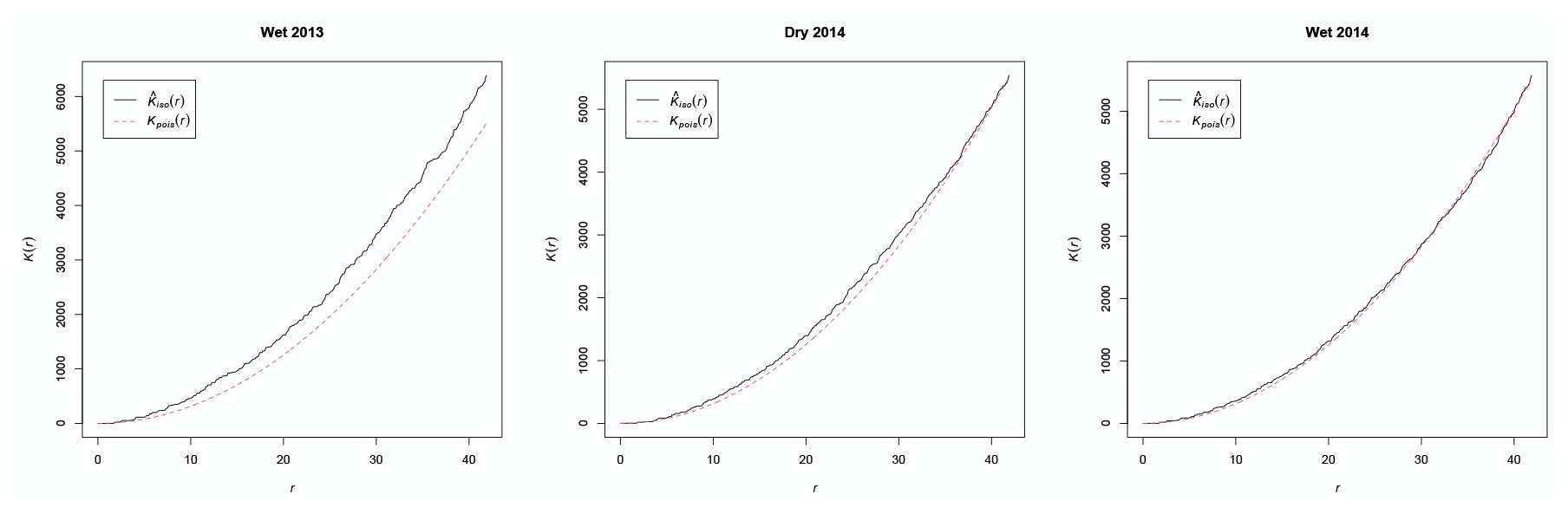
Results of Ripley’s *K* test, which quantifies the cumulative average number of trees *K* lying within a distance *r* (in meters) of a typical data point. 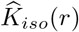 corresponds to the observed data for each season, while *K*_*pois*_(*r*) is the theoretical curve assuming an inhomogenous Poisson point process. As can be seen, the fit between the theoretical and observed curves is better for both 2014 seasons than for the wet season of 2013, when there is some evidence of a higher intensity of clustering. The isotropic correction for edge effects and a normalization by intensity was applied to 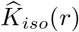.. Figures generated using the *spatstat* package for R [R Core Team, 2021, Baddeley et al., 2015].

The use of a model with different intensities for each species in the Poisson model was justified by comparing its fit to the observed data in the 1-ha plot to that of an alternative model which assumed only spatial inhomogeneity but no differences between species. Table 1 shows the results of these tests for each season.

**Table 1:** Results of the likelihood ratio test copmaring, for each season and year combination, the fit of a Poisson model assuming the same intensity for all trees (Poisson) with another Poisson model assuming different a different intensity for each species (Poisson + species). In all cases, deviances are significantly different with a P<0.0001 (*) and the Akaike information criteria (AIC) is lower for the Poisson model with different intensities for each species, despite the larger number of parameters (6 for Poisson, 15 for Poisson + species). All analyses were performed using the *spatstat* package for *R* [R Core Team, 2021, Baddeley et al., 2015].

| Season, year | Model | Deviance | AIC |
| --- | --- | --- | --- |
| Wet, 2013 | Poisson |  | 1554 |
|  | Poisson + species | 187.2* | 1385 |
| Dry, 2014 | Poisson |  | 2525 |
|  | Poisson + species | 263.2* | 2279 |
| Wet, 2014 | Poisson |  | 2575 |
|  | Poisson + species | 382.8* | 2210 |

